# Beyond copy number: The regulatory architecture of mitochondrial DNA gene expression

**DOI:** 10.64898/2026.05.27.728277

**Authors:** Parisa Riahi, Sharwary Raghupathy, Bryan Le, Sol Taylor-Brill, Dylan Taylor, Rajiv McCoy, Shweta Ramdas, Arslan A. Zaidi

**Author notes:** Correspondence to A.A.Z. Contributed equally (Parisa Riahi, Sharwary Raghupathy, Bryan Le).

## Abstract

Mitochondrial DNA (mtDNA) copy number is widely used as a biomarker for mitochondrial function and disease risk, yet its relationship to mtDNA gene expression – one important functional output – remains poorly understood. Prior studies examining this relationship have largely relied on heterogeneous tissue samples, where confounding by cell-type composition obscures the underlying biology. We rigorously tested this relationship in lymphoblastoid cell lines (LCLs), where we find no correlation between mtDNA copy number and gene expression across 731 individuals, and minimal association across 49 GTEx tissues except whole blood. Using population genetic modeling of heteroplasmy drift between DNA and RNA, we estimate that effectively ≈50 out of 813 mtDNA templates are transcriptionally active in LCLs, indicating low mtDNA accessibility. This confirms, using an independent method and a different cell type, previous observations in HeLa cells, where mtDNA is largely compacted into nucleoids. Together, our results demonstrate that mtDNA copy number and expression are largely decoupled, and that above a certain rate-limiting threshold, mtDNA accessibility — rather than absolute copy number — is the more relevant quantity for explaining inter-individual differences in gene expression. This challenges the interpretation of mtDNA copy number as a proxy for mitochondrial transcriptional output and highlights the need for a more mechanistic understanding of mtDNA copy number associations with disease-relevant traits. *Cis*- and *trans*-eQTL mapping further reveals that genetic regulation of mtDNA gene expression operates primarily through post-transcriptional mechanisms rather than transcription initiation, yet despite the high inter-individual variance in mtDNA gene expression, genetic variation underlying mtDNA regulation appears to be under strong selective constraint.

## Introduction

Mitochondria are essential for cellular function. They are best known for their role in the generation of ATP through oxidative phosphorylation (OXPHOS) but also play important roles in apoptosis, nucleotide synthesis, cell signaling, and nutrient metabolism [1–3]. Therefore, it is not surprising that mitochondrial disorders are multisystemic and mitochondrial dysfunction underlies many common diseases such as diabetes [4], cardiovascular disease [5], and cancer [6]. Gold standard metrics of mitochondrial (dys)function such as oxygen consumption, reactive oxygen species (ROS) concentration, and mitochondrial membrane potential cannot be measured at scale. Thus, there is substantial interest in the field in identifying non-invasive, high throughput biomarkers of mitochondrial (dys)function.

Mitochondrial DNA (mtDNA) copy number (mtCN), the average number of copies of mtDNA per cell, has emerged as one such promising biomarker because of the ease with which it can be estimated at scale from DNA sequencing data [7]. MtCN is known to correlate with mitochondrial activity within individuals, such that metabolically active tissues such as cardiac and skeletal muscle cells carry more copies of mtDNA than tissues that are less active [8]. MtCN is also correlated with mtDNA gene expression, consistent with the demand for proteins needed for ATP production [9, 10]. MtDNA transcription is polycistronic and is not regulated by canonical enhancer elements as in the case of nuclear genes. As such, mtDNA transcription is thought to be regulated by varying mtCN, which therefore has the potential to be predictive of mtDNA gene expression and mitochondrial activity across individuals. However, the empirical support for the relationship between mtCN and mtDNA gene expression comes from analyses in heterogeneous tissues [8, 11], where biological factors that vary across individuals (e.g. cell composition) can induce transcriptome-wide correlations that are not direct effects of mtCN. Because of this, the causal relationship between mtCN and mtDNA gene expression and, therefore, the predictive utility of mtCN remains in question.

To address this, we studied the regulatory landscape of mtDNA gene expression and the predictive ability of mtDNA copy number with whole genome and transcriptome sequencing data from lymphoblastoid cell lines (LCLs) from 731 individuals who were part of the 1000 Genomes Project. The use of cell lines limits confounding due to cell heterogeneity. We test for an association between mtCN and mtDNA gene expression through an analytical framework grounded in causal reasoning that minimizes different sources of biases that can distort this relationship. We also developed a method that uses RNA and DNA sequence data to estimate the transcriptional bottleneck – the effective number of mtDNA copies under active transcription – complementing absolute copy number measurements by providing insight into mtDNA accessibility and template usage. Finally, we carried out *cis-*QTL analyses to study the genetic basis of mtDNA gene expression. Our findings reveal that mtCN is a surprisingly weak predictor of mtDNA gene expression, and that mtDNA accessibility, post-transcriptional regulation, and nuclear genetic control are likely more important in explaining inter-individual variation in mtDNA gene expression.

## Results

### Little evidence of direct effect of mtCN on mtDNA gene expression

We analyzed published data from the Taylor *et al*. (2024) [12], who sequenced the transcriptome of lymphoblastoid cells lines (LCLs) derived from the peripheral blood mononuclear cells (PBMCs) of 731 donors from the 1000 Genomes Project. RNA sequencing was performed via polyA enrichment, which allowed us to obtain raw counts of the 12 mtDNA protein-coding genes that are polyadenylated post-transcriptionally [13, 14]. We estimated mtCN from high coverage (median depth on mtDNA = 14, 601×) whole genome sequences from the same LCLs [15] (Methods, Data S1). MtDNA copy number estimated from the high-coverage data was slightly lower than that estimated from low-coverage (median depth = 3, 472×) data from the same samples [16] (Fig. S1), potentially reflecting mtDNA loss during prolonged LCL culture. However, the estimates were strongly correlated (Fig. S1), demonstrating that inter-individual variation in copy number remains stable across cell passages.

The association we seek to estimate includes both the direct effect of mtCN on mtGE and effects of shared regulators such as TFAM, which influences both mtCN and mtGE independently [17]. However, to estimate this accurately, we need to account for spurious correlations through other mechanisms, which we illustrate using a directed acyclic graph (DAG) in Fig. 1A. First, biological confounders such as age, sex, and cell composition affect both mtCN and gene expression. Technical confounders – variables affecting sequencing experiments such as batch effects and library size – can also induce gene expression correlations, though they are less of a concern here as RNA and DNA were sequenced independently. Both types of confounders can be addressed either by conditioning on measured variables, or more commonly, through surrogate variables (e.g. PEER factors or principal components) derived from expression counts. Second, standard library size normalization, which is equivalent to conditioning on the total number of reads – a collider – could introduce an underappreciated source of bias. Because mtDNA-encoded genes exhibit high mean expression and variance (Fig. S2A), they contribute disproportionately to total read count. Thus, conditioning on total read count can induce spurious transcriptome-wide correlations with mtDNA gene expression, potentially distorting the mtCN-mtGE association in unpredictable ways. To avoid this, we exclude mtDNA genes from the normalization step. Third, including surrogate variables as covariates – while intended to remove confounding — can overcorrect by absorbing mtDNA gene expression variation if mtDNA genes are included in their calculation, attenuating the estimated mtCN-mtGE association. We demonstrate this empirically by showing that the top principal components computed from gene expression (ePCs) (Methods) capture substantial mtDNA gene expression variation when mtDNA genes are included in the PCA, but not when they are excluded (Fig. S3A). PEER factors previously computed by GTEx [18] and MAGE [12] were similarly correlated with mtDNA expression (Fig. S3C). We therefore exclude mtDNA genes from ePC calculation as well. Overall, our approach addresses potential confounding and collider biases while preserving expression variation relevant to mitochondrial function. To detect any residual bias, we use pseudogenes and long non-coding RNA (lncRNA), which were excluded from normalization and ePC calculation, as negative controls.

**Figure 1:**
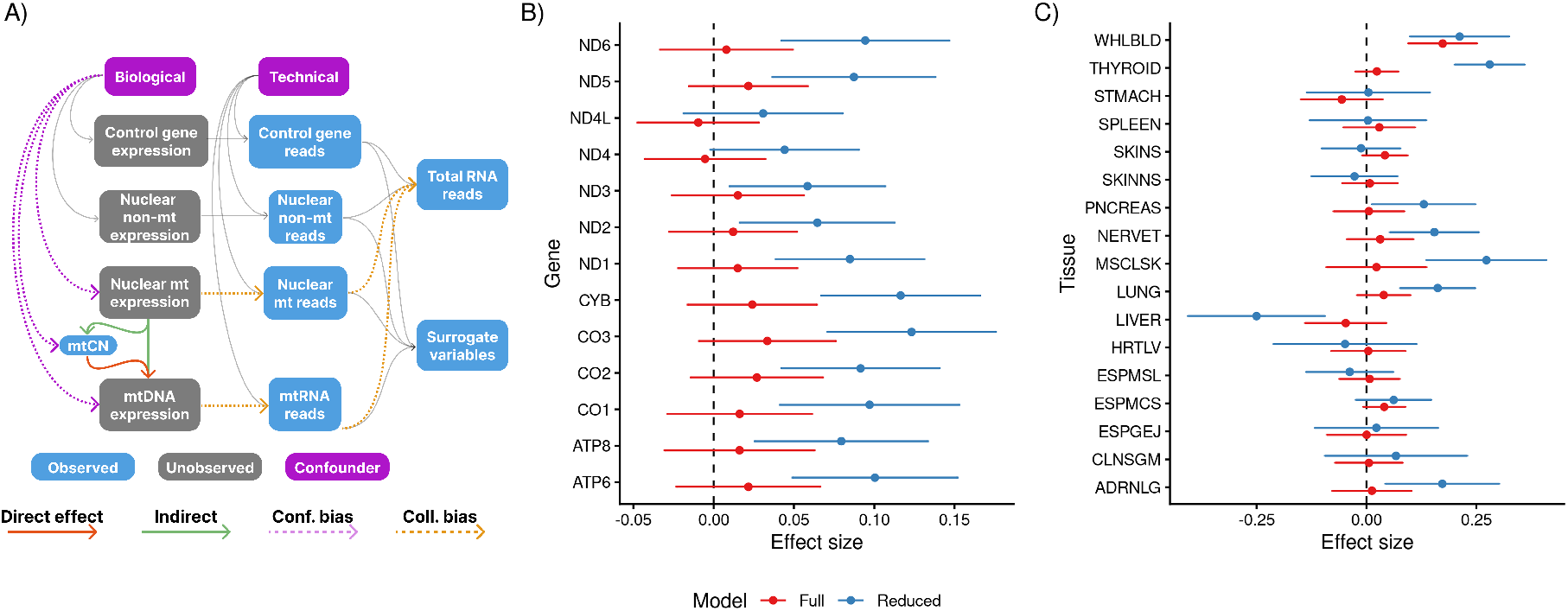
(A) Directed acyclic graph (DAG) representing a generative model underlying the association between mtCN and mtGE. Observed and unobserved variables are shown in grey and blue boxes, respectively, and confounders are shown in purple. Causal paths are depicted as arrows with direct effects of interest in red, indirect effects in green, and biasing paths because of confounding and collider bias in pink and orange dotted lines, respectively. Only biases relevant to the exposure and outcome of interest are highlighted. (B) Effect size of mtCN on expression of mtDNA genes in LCLs. (C) Effect size of mtCN on mtDNA gene expression in different tissue types. Only the effect on *MT-CO1* is shown here and the effect on other genes is shown in Fig. S6. Reduced model only includes sex, age, age^2^, batch, and gPCs as covariates. Full model includes covariates in reduced model as well as ePCs.

We tested for an association between mtCN and the gene expression of 13 protein-coding mtDNA genes using a linear model with batch, sex, first five genetic PCs (gPCs), and ePCs computed on normalized expression counts as fixed effects to account for genetic ancestry and other confounders (Methods). In LCLs, the expression of most mtDNA genes was positively correlated with mtCN but this correlation attenuates to non-significance upon inclusion of ePCs (Fig. 1B), suggesting that the raw association is driven by confounders. Pseudogenes and long non-coding RNA (lncRNA) associations were inflated without ePCs (*λ* = 1.2 for pseudogenes and *λ* = 1.5 for lncRNA) and well-calibrated (*λ* ≈ 1 for both sets, Fig. S4) upon their inclusion as covariates. We repeated this analysis in 17 GTEx tissues with sufficient sample sizes (Methods). Without ePCs, the association between mtCN and mtDNA gene expression was significant in several tissues but were inconsistent in direction — positive in whole blood, negative in liver (Fig. 1C) — and mtCN showed transcriptome-wide inflation in its correlation with gene expression (Fig. S5). Inclusion of ePCs corrected this inflation (Fig. S5), leaving only whole blood with a significant mtCN-mtGE association (Fig. 1C). The association in whole blood is consistent with Yang *et al*. [11], but our results are better calibrated (*λ*_pseudogene_ = 0.99, *λ*_lncRNA_ = 1.1, Fig. S5). We examined the correlation between mtDNA gene expression and genetically predicted mtCN (pmtCN), which should be free of confounding, provided the mtCN-associated variants and their effects are not biased by cell composition. We found no significant associations in any tissue or cell type (Fig. S7), with well-calibrated negative control associations (Fig. S8). Overall, results from both observed and genetically predicted mtCN suggest the absence of a consistent effect across tissues.

The residual association in whole blood likely reflects platelet contamination rather than a genuine within-cell regulatory relationship. Platelets, which lack nuclear DNA but retain mtDNA, contribute disproportionately to blood-derived mtCN estimates [19] and also exhibit high mtDNA expression [20]. As a result, platelet abundance induces a spurious correlation between bulk mtCN and mtDNA gene expression. This confounding effect is expected to be non-linear in a way that ePCs – which capture linear axes of variation – would fail to correct. Replicating this association in a dataset with directly measured platelet counts will be important for determining whether it reflects a within-cell relationship between mtCN and mtDNA gene expression. Nevertheless, the association in whole blood is weak (*r*^2^ = 0.02), even if it reflects a genuine within-cell relationship. Altogether, inter-individual variation in mtDNA copy number does not meaningfully explain mtDNA expression variation.

### An mtDNA transcriptional bottleneck in LCLs

The observation that mtCN is uninformative about mtDNA gene expression in most tissues is consistent with evidence that mtDNA accessibility is likely more important for regulation of mtDNA genes than absolute copy number [21]. A recent study showed through single molecule sequencing that despite there being a large number of mtDNA molecules in HeLa cells, most are compacted into inaccessible nucleoids [21]. This observation implies a ‘transcriptional bottleneck’ where most mtRNA transcripts would arise from a small number of accessible mtDNA molecules. We developed a population genetic model to test this in LCLs (Fig. 2) (Methods). Briefly, we discovered heteroplasmies in the mtDNA sequence data from the high-coverage 1000 Genomes sequencing [15] and mtRNA sequence data from MAGE [12] and estimated the amount of drift experienced by the heteroplasmies during transcription (*ψ*). Assuming this drift occurs in a single event, we can estimate the effective number of mtDNA that were transcribed – the transcriptional bottleneck – as the inverse of this drift parameter 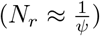. A large drift parameter implies fewer transcriptionally active copies and therefore a tighter bottleneck, and vice versa.

**Figure 2:**
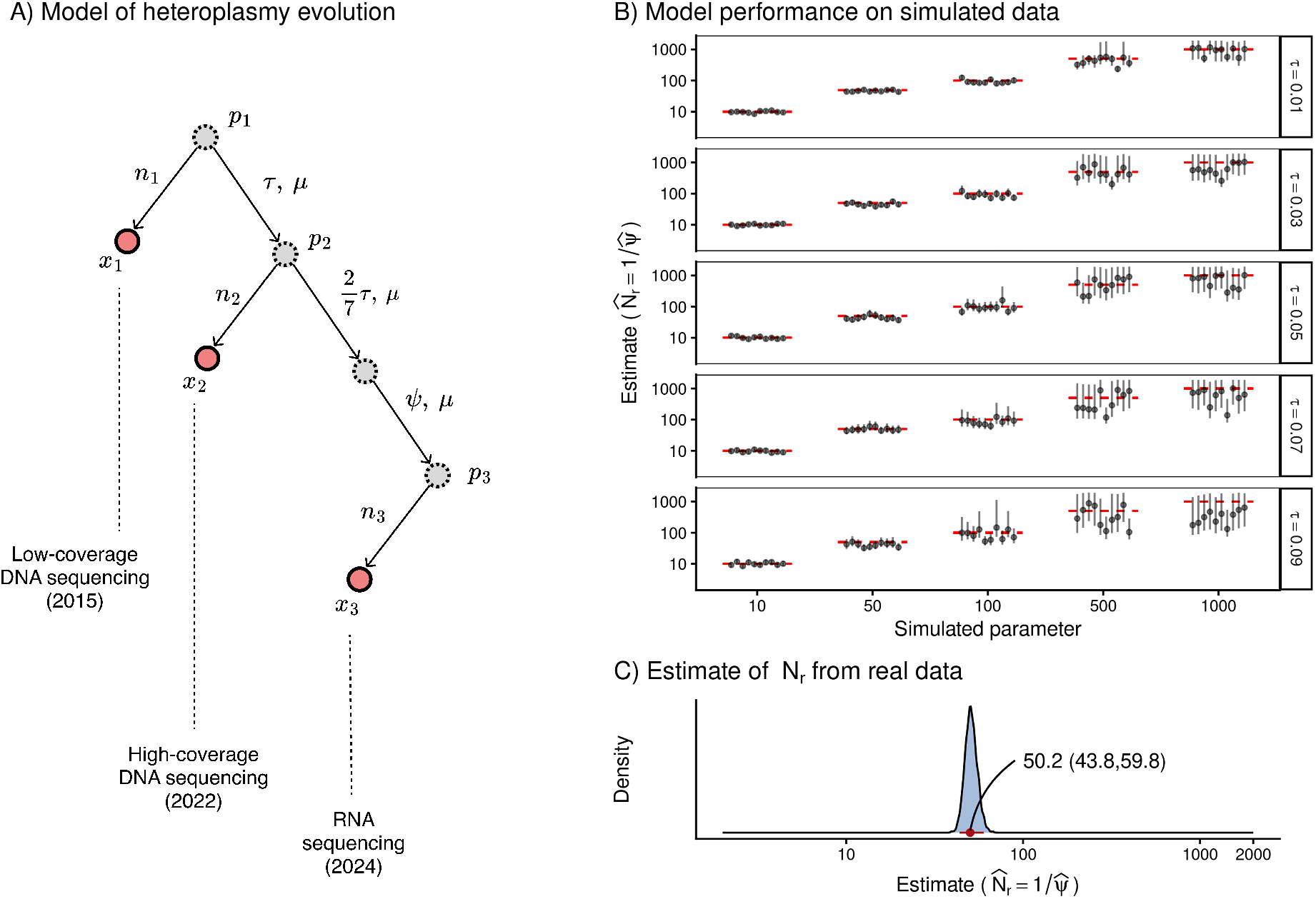
(A) Model of evolution in heteroplasmy frequencies used to estimate drift due to cellular/mtDNA turnover (*τ*) and transcriptional bottleneck 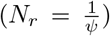. Observed counts (*x*_*i*_) are shown as red circles while latent allele frequencies (*p*_*i*_) are shown in grey circles. *n*_*i*_ refers to read depth at the *i*^*th*^ heteroplasmy while *µ* represents the mutation rate. (B) The model yields unbiased estimates of *N*_*r*_ in simulations. Red dashed lines represent the true simulated value of *N*_*r*_ and the black points and vertical bars represent the MAP estimates and 95% CrIs, respectively. Results are shown for 10 different replicates for a range of simulated values of *N*_*r*_ and *τ*. (C) The inferred posterior distribution of *N*_*r*_. The x-axis is scaled to represent a log-uniform prior in that range.

One challenge here is that the DNA, which was sequenced in 2022 [15] does not represent the parent mtDNA population from which the RNA was transcribed in 2024 [12]. Failure to account for any drift accumulated in the mtDNA during this time period due to mtDNA/cell turnover would inflate drift estimates, leading to an underestimate of the transcriptional bottleneck size. We account for this by leveraging heteroplasmy data from an additional time point: the low-coverage sequencing of the same samples carried out in 2015 [16]. We make the assumption that mtDNA turnover is linear in time and therefore the drift experienced by heteroplasmies between 2022 and 2024 (*τ*) is ≈ 2*/*7× that of the drift accrued between 2015 and 2022 (Fig. 2A). We can estimate, and therefore account for, this drift from the observed change in heteroplasmy frequencies between the two DNA sequences. We also account for any variance in heteroplasmy frequency due to mutations and site-specific sequencing depth (Methods).

We use a Bayesian approach to inference (Methods). Our model accurately recapitulates heteroplasmy evolution under the Wright-Fisher model (Fig. S9) and yields unbiased estimates of the bottleneck parameter in simulations (Fig. 2B). Credible intervals (CrIs) can be wider, however, for large bottleneck sizes (e.g. *N*_*r*_ ≥ 500), especially if there is substantial drift due to mtDNA turnover. This is because there is very little drift in large populations, requiring more sites to estimate accurately. Nevertheless, a lack of precision in this regime does not preclude us from confidently detecting small bottlenecks. We applied this model to 418 single nucleotide heteroplasmies that were detected with a minor allele fraction of > 0.01 at a depth of > 1,000x in the low-coverage DNA sequence (tree root in Fig. 2A, Data S2). Our analysis yielded a maximum a posteriori (MAP) estimate of 50.2 (95% CrI: 43.8 – 59.8) effective units of mtDNA templates/cell (Fig. 2C). Given that the mean copy number in the same samples is 813 copies/cell, our estimate implies that relatively few (≈ 6%) of mtDNA molecules in LCLs undergo transcription, confirming the observation in HeLa cells, where most mtDNA is compacted into inaccessible nucleoids [21].

### *Cis*- and *trans*-eQTLs converge on post-transcriptional regulation of mtDNA expression

The absence of an effect of mtCN in LCLs raises the question of whether inter-individual variation in mtDNA expression is instead regulated by genetic variants. To answer this, we performed eQTL analysis for the expression of 13 protein-coding mtDNA genes. We first performed *cis*-eQTL analysis with 411 mtDNA SNPs with MAF > 0.01 that passed QC (Methods), identifying 47 that passed a stringent threshold of 5 × 10^−9^ (Methods, Data S3). We fine-mapped these associations with `SuSiE` [22] to 12 credible sets across all genes (Fig. 3A, Data S4), suggesting that most associations are driven by high linkage disequilibrium (LD) in the mtDNA. Indeed, all associations within a credible set were in high LD (Fig. S10) and often represented haplotypes within broadly-defined mtDNA haplogroups (Fig. S11). For example, *ATP8* had the largest credible set with 15 variants tagging the L0 haplogroup, which is most commonly found among people of African ancestry (Fig. S11). The effect estimates of variants associated with mtDNA expression in LCLs were largely concordant with their estimates in whole blood (Fig. 3B), suggesting that the genetic architecture of mtDNA expression might be shared across tissues.

**Figure 3:**
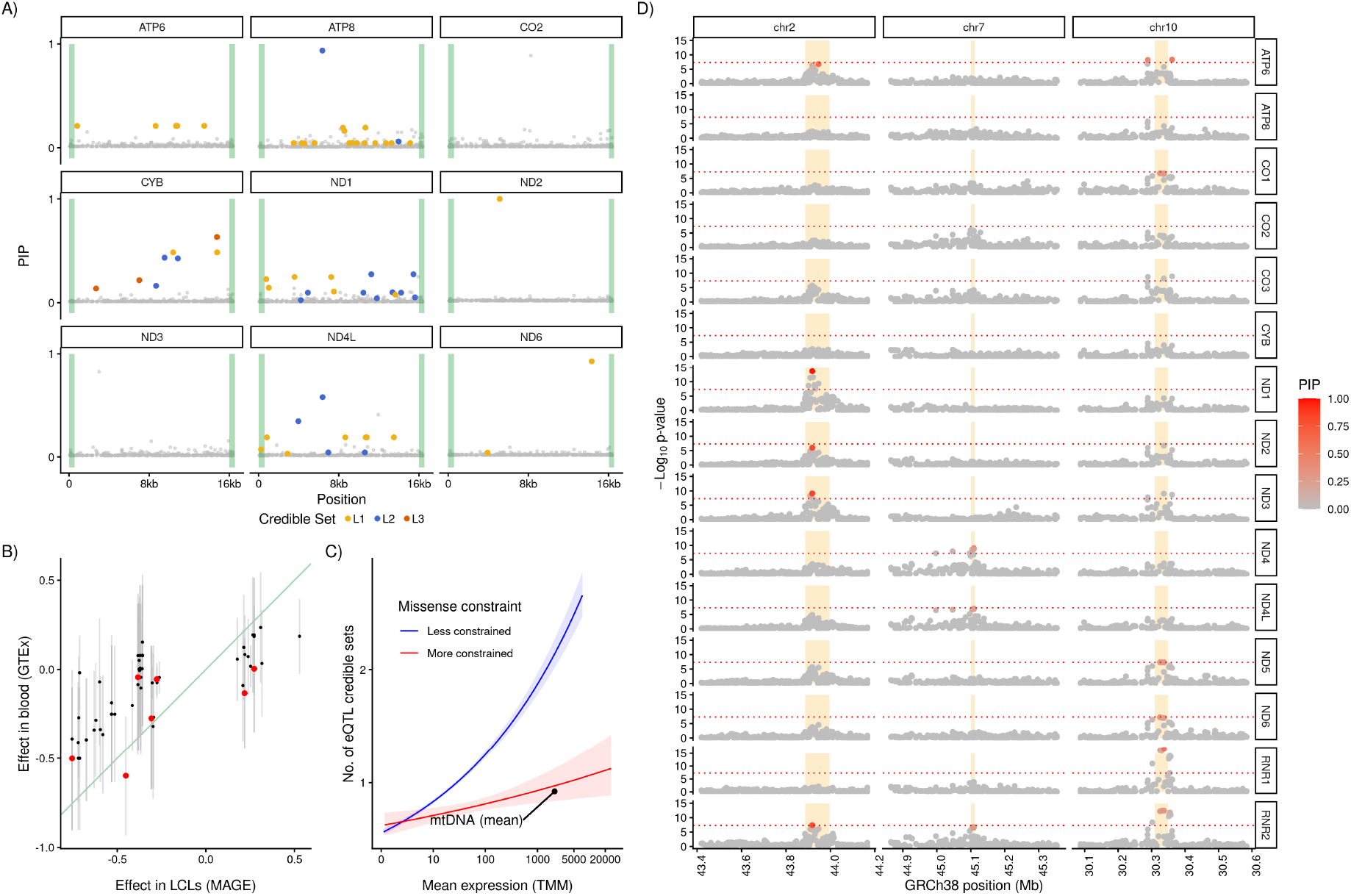
Expression QTL (eQTL) analysis of mtDNA gene expression in MAGE. (A) *Cis*-eQTL results where the position on the mitochondrial genome (rCRS) is shown on the x-axis and the posterior inclusion probability (PIP) is shown on the y-axis. Colored points represent variants in fine-mapped credible sets and the shaded green segment represents the D-loop region. (B) Comparison of effect sizes of associated *cis*-eQTLs between LCLs (x-axis) and whole blood (y-axis). The red points are the variants with the highest PIP within each credible set identified in MAGE that were also present in GTEx. The green line represents the *y* = *x* line. (C) Number of *cis*-eQTL credible sets in the nuclear genome (y-axis) modeled as a function of mean expression level (x-axis) and missense constraint (color) using a quasi-Poisson model (Methods). The number of credible sets identified per gene in the mtDNA is shown for comparison. (D) *Trans*-eQTL results for mtDNA gene expression in MAGE. Only variants in a 1Mb region around the three most significant associations are shown. The y-axis represents the -log_10_(p-value) of the association and the color represents the PIP of each variant. The shaded yellow segment represents the coding region of the gene harboring the lead SNP.

Most fine-mapped variants resided outside the D-loop region (Fig. 3A). Of the credible sets, only one (for *ND4L*) contained a variant in the D-loop (m.247:G>A) within the origin of heavy strand replication (OHR) upstream of conserved sequence blocks 2 and 3. The OHR serves as the initiation site for mtDNA replication and transcription and this variant might affect DNA binding of proteins that are components of the transcriptional machinery. This variant was significantly associated with *ND4L* and *ATP6* expression and was nominally significant (*p <* 1 × 10^−4^) for *ND1, ND2, ND3*, and *ATP8* expression. All other fine-mapped variants fell outside the D-loop and spanned the length of mtDNA. This is surprising because both the light- and heavy-strand promoters reside within the D-loop region, suggesting that steady state variation in mtDNA gene expression in LCLs is likely driven by post-transcriptional processing and mRNA stability as opposed to transcription initiation, in agreement with previous findings [23–26].

We observed fewer credible sets for the *cis*-eQTLs than expected given the high mean and variance in expression of mtDNA genes in LCLs (Fig. 3C). We hypothesized this might be because of selective constraint on mtDNA gene expression since highly expressed nuclear genes under selective constraint tend to be depleted in eQTLs [12, 27] (Fig. 3C). To evaluate this, we downloaded constraint scores for nuclear and mtDNA genes from gnomAD v4.1.0 [28, 29]. We chose the 90% upper bound of the ratio of the observed vs expected number of missense variants observed in gnomAD to measure constraint and labeled the 10% of nuclear genes with the lowest scores as highly constrained and the bottom 90% as less constrained. We show that mtDNA genes are highly constrained, with missense constraint scores that are even more extreme than the most constrained genes in the nuclear genome (Fig. S13). Thus, genetic regulation of mtDNA gene expression appears to be highly constrained, despite its high variability, across both individuals and tissues.

The role of post-transcriptional processing and mtRNA stability in regulating mtDNA gene expression was also evident in the results of *trans*-eQTL analysis, which we performed with ≈ 1*M* variants across the nuclear genome (Methods). We detected three associations that passed a study-wide threshold of *p <* 5 × 10^−9^ and that were shared by multiple mtDNA genes (Fig. 3D, Fig. S12, Data S5). The lead SNPs resided in nuclear genes known to be involved in mtRNA processing. The lead SNP on chromosome 2 resided in the *LRPPRC* gene, which binds to the poly(A) tails of mtDNA transcripts to regulate their stability [30]. The lead SNP on chromosome 7 resided in the *TBRG4*, also known as *FASTKD4*, which binds to mtDNA transcripts and modulates their half-lives [31]. The lead SNP of the chromosome 10 association lay in the *MTPAP* gene, the polymerase responsible for creating poly(A) tails on mtDNA transcripts. The fine-mapped variants with the highest PIP in each gene are known *cis*-eQTLs for these genes in multiple GTEx tissues, suggesting that the effects of these eQTLs on mtDNA gene expression are likely mediated by regulating the transcription of nuclear-encoded genes. Altogether, our findings indicate that both local and distal genetic variation converge on post-transcriptional mechanisms to regulate mtDNA transcript abundance, in accordance with previous studies [23, 24, 26].

## Discussion

The factors contributing to inter-individual variation in mtDNA gene expression are not well-studied. A common hypothesis is that mtCN, which generally correlates with mtDNA gene expression levels, at least across tissues [8], drives this difference. This makes sense given that mtDNA does not carry canonical enhancer elements and therefore, one way to regulate mtDNA gene expression is by regulating mtDNA copy number. Indeed, tissues with higher energetic demand (e.g. skeletal muscles) exhibit higher expression and copy number of mtDNA compared to tissues with lower energetic demand (e.g. blood) [8]. However, the observed cross-tissue correlations between mtCN and gene expression may not translate to a correlation across individuals within the same tissue or cell type. Correlations within heterogeneous tissues (e.g. whole blood [11]) are also susceptible to confounding by cell heterogeneity. To address this, we investigated the relationship between mtCN and mtDNA gene expression in lymphoblastoid cell lines [12]. Our findings shed light on a complex regulatory architecture of mtDNA gene expression.

We did not observe a significant association between mtDNA copy number and expression or between genetically predicted mtDNA copy number and expression in LCLs. Thus, there is little empirical evidence that mtDNA copy number is predictive of mtDNA expression across individuals, conditioning on cell heterogeneity. We also did not find a significant relationship between mtDNA copy number and expression in any tissue in GTEx except for whole blood. Since the GTEx data did not have cell composition information, we cannot fully rule out the confounding effects of cell composition in driving the association in blood. It might represent a real effect of mtDNA copy number on gene expression but this effect is still small as mtDNA explains ≈ 2% of the variance in mtDNA gene expression. Blood has the lowest average mtDNA copy number and gene expression of all GTEx tissues [32]. Thus, the observed association in blood might be indicative of a threshold effect of mtCN on mtDNA gene expression. Some mtDNA templates are, obviously, needed for transcription, but as long as there are enough copies, the absolute number might be less important. This is consistent with a recent study in HeLa cells that showed that despite there being a large number of mtDNA copies, only a fraction are accessible [21]. This would suggest that as long as cells have enough templates, the number of accessible copies of mtDNA and not the absolute number is likely important for regulating mtDNA gene expression.

We studied mtDNA accessibility in LCLs by developing a statistical method grounded in population genetics. The key insight is that transcription samples from the mtDNA pool, and this sampling introduces drift in heteroplasmy frequencies between mtDNA and mtRNA. The magnitude of this drift is informative about how many copies were effectively sampled – the transcriptional bottleneck (*N*_*r*_). In LCLs, we estimate that ≈ 50 effective copies of mtDNA underwent transcription. Given an average copy number of 813, our estimate of *N*_*r*_ implies reduced accessibility (≈ 6%) of mtDNA in LCLs, confirming, in an independent cell type, the limited mtDNA accessibility observed in HeLa cells [21]. However, note that *N*_*r*_ is different from direct measures of accessibility in that the latter is a snapshot while the former provides a marginal estimate over time. But this duration is not expected to be long given that the half-life of mtRNA is on the order of hours [33, 34]. Beyond LCLs, this framework could in principle be applied to estimate transcriptional accessibility across cell types and physiological conditions where matched mtDNA and mtRNA sequencing data are available.

Finally, we carried out eQTL analysis to study the contribution of genetic variation to mtDNA gene expression. We discovered several *cis-* and *trans-*eQTL associations. All except one of the *cis-*eQTLs resided outside the D-loop, where most of the transcriptional initiation occurs. The *trans-*eQTLs resided in genes involved in post-transcriptional processing and mtRNA stability. Altogether, this suggests that genetic regulation of mtDNA gene expression in LCLs operates primarily through transcript stability and post-processing rather than initiation. We also showed that most fine-mapped eQTLs tag mtDNA haplogroups, indicating that between-population genetic variance contributes, at least partly, to mtDNA gene expression. Despite this, genetic regulation of mtDNA gene expression is under strong selective constraint.

Our results do not imply that there is no causal relationship between mtDNA and gene expression. Some template is obviously needed for mtDNA transcription and targeted depletion of mtDNA results in mtRNA depletion [35]. Knockout of genes essential for mtDNA replication such as *POLG, POLG2*, and *TK2* result in loss of mtDNA and therefore, of mtDNA gene expression and mitochondrial activity [36, 37]. Similarly, recent knockdown of *POLG* and *POLG2* with perturb-seq also found mtDNA genes to be among the top downregulated genes in both cell lines studied [38]. But this relationship is likely context-dependent and non-linear. For example, *TK2* deficiency depletes mtDNA in both brain and heart, yet expression loss occurs only in brain — pointing to tissue-specific compensatory mechanisms [39]. Similarly, moderate over-expression of *TFAM* can result in an increase in mtDNA copy number without a concomitant increase in gene expression [7] but higher *TFAM* levels can reduce expression by compacting mtDNA into nucleoids [21]. This points to a non-linear relationship between mtDNA copy number and gene expression that should vary by cell type. The residual association we observe in whole blood – the tissue with the lowest average mtCN – is consistent with this: absolute copy number may matter most when templates are limiting.

Our results also do not imply that mtDNA copy number is uninformative about disease or that it has no value as a biomarker given that it is widely associated with many diseases [7, 40, 41]. But these associations, even if they are causal, reflect the marginal effect of many processes besides mtDNA transcription that link mtDNA copy number to mitochondrial (dys)function. These effects are almost certainly not the same across individuals and likely depend on their genetic makeup, health, and environmental context. Thus, for mtDNA copy number to be a useful biomarker or to understand its relationship to human health in general, a mechanistic understanding of these processes is needed.

## Materials and Methods

### Quantifying gene expression and mtCN in MAGE

We used the Multi-ancestry Analysis of Gene Expression (MAGE) resource to study the regulation of mtDNA gene expression in lymphoblastoid cell lines (LCLs). MAGE comprises RNA-seq data from LCLs derived from peripheral blood mononuclear cells (PBMCs) of 731 individuals across geographically diverse human populations [12]. Gene expression was quantified previously by Taylor *et al*. using `Salmon` (version 1.5.2) [42], which uses an alignment-free approach to estimate transcript-level pseudocounts. These were summed to gene-level counts using the Gencode v38 transcript annotation. We used `edgeR` [43] to normalize read counts with respect to gene length and library size. To avoid introducing compositional bias, we computed the normalization factor only from a set of protein-coding genes that excludes genes known to be involved in mitochondrial function. To do this, we selected a list of 13,271 genes that are (i) annotated as protein coding in Gencode v38, (ii) not listed in Mitocarta 3.0 [44], and (iii) have TPM of > 0.1 and read count > 6 in at least 20% of individuals. This factor was then applied to all genes with the `cpm()` function in `edgeR` to compute TMM values, which were then used to represent gene expression in all downstream analyses. We computed ePCs using the `prcomp()` function in R [45] on the normalized expression of the set of 13,271 genes mentioned above to capture axes of global expression variation driven by biological or technical confounders. We included 100 ePCs in the model as fixed effects.

MtDNA copy number (mtCN) was estimated from the high-coverage [15] and low-coverage [16] whole genome sequencing data of the 1000 Genomes Project. To do this, we ran `samtools idxstats` (version 1.16.1) [46] on the CRAM files to get the total number of mapped reads to the autosome and mitochondrial genomes. Then mtCN was computed as the ratio of the average mitochondrial depth to the average nuclear depth × 2. We computed polygenic scores for mtCN with effect sizes of 110 independent SNPs identified in Longchamps *et al*. [47] using `plink2 --score` [48, 49].

### GTEx analysis

We analyzed the relationship between mtCN and gene expression in 17 tissues from GTEx v8 [18] using mtCN measured by Rath *et al*. (2024) [32]. We normalized gene expression and computed ePCs with the same approach as outlined in Fig. 1 separately in each tissue. We fit a model separately in each tissue with the same set of covariates as used by the GTEx eQTL analysis [18], but using ePCs instead of PEER factors.

### mtDNA variant Calling

We called mtDNA and mtRNA variants for each sample separately using a modified version of the published gnomAD pipeline [50] from the whole genome and transcriptome alignment files generated previously for the 1000 Genomes [12, 15] and GTEx v8 [18]. The artificial start and end points of the revised Cambridge Reference Sequence (rCRS) of the human mtDNA, which is circular, can result in a small loss of reads spanning the D-loop region. To account for this, the pipeline calls variants twice, once from reads aligning to the original reference and then from a shifted reference alignment where the start position correspond to 8000bp of the rCRS. For each alignment, only properly paired reads with both mates aligning to chrM were retained to reduce artifacts. The alignment was then converted to FASTQ using `Picard SamToFastq` [51] and re-aligned to the mitochondrial reference using `BWA-MEM` [52]. For RNA analysis, secondary and supplementary alignments were removed to avoid double-counting of multi-mapping reads. Alignments were merged with original metadata using `Picard MergeBamAlignment` [51]. Duplicated reads were removed prior to variant calling.

Variant calling was done using `GATK mutect2` [53]. Multi-nucleotide polymorphisms were disabled (–max-mnp-distance 0) to report single nucleotide and indel events independently, and the maximum number of reads per alignment start was capped (–max-reads-per-alignment-start 75) to reduce artifacts arising from read stacking. Variants with base quality < 30 and/or map quality < 20 were removed using `GATK FilterMutectCalls` [53]. The resulting variants from both alignments are then merged into a single VCF for each sample and duplicate records removed. Thus, Each VCF contains the fraction of alternate allele carrying reads representing the heteroplasmy fraction of the ALT allele (HF) in that individual’s tissue. We reduced the number of low-confidence heteroplasmies potentially attributed to sequencing error by applying a heteroplasmy threshold of 0.01, which excludes variants with HF < 0.01 or > 0.99 in each VCF. We further filtered our variants present exclusively in RNA to avoid effects of post-transcriptional modification or RNA editing. Positions present in DNA but absent in RNA were retained as these might be alleles that drifted to fixation or loss as a result of mtDNA turnover or transcriptional bottleneck. Variants failing the gnomAD quality filters in either DNA or RNA, and problematic, artifact-prone sites (66–71, 300–317, 514–523, 316, 3105–3109, 12418–12425, 16180–16193) were excluded from downstream analyses.

### Estimating the transcriptional bottleneck

We used heteroplasmy data to estimate the average number of copies of mtDNA per cell that were transcribed through an approach based in population genetics. Intuitively, heteroplasmies present in mtDNA will experience more variance, i.e., drift, if fewer molecules are sampled during transcription compared to the situation where most of the mtDNA is transcribed. Thus, in principle, we could estimate the drift, and therefore, the ‘transcriptional bottleneck’ from the variance in heteroplasmy fractions between the mtDNA and RNA populations. This is more complicated in practice because the RNA sequencing was carried out in 2024 while the high-coverage DNA was sequenced in 2022. Thus, the RNA was transcribed from an mtDNA pool that was not directly sequenced and any drift in heteroplasmy frequency between 2022 and 2024 due to cell passage or mtDNA turnover will result in an over-estimation of drift during transcription and, therefore, an underestimation of the transcriptional bottleneck. To account for this, we leveraged the low-coverage DNA sequencing of the 1000 Genomes samples from 2015. Assuming the drift due to mtDNA turnover is linear in time, the drift between the mtDNA pool sequenced in 2022 and the mtDNA pool from which the RNA was transcribed is 2/7^th^ of the drift experienced between 2015 and 2022. We use a Bayesian framework to account for this extra drift as well as any variance due to mutation and uneven read depth.

We used a hidden Markov model (HMM) similar to models used to study the germline bottleneck [54–56] to estimate the transcriptional bottleneck (*N*_*r*_). The latent allele frequencies at each node of the tree (Fig. 2A) constitute hidden states while the observed allele counts are emissions. For the *i*^*th*^ heteroplasmy discovered in the low-coverage mtDNA sequence, let 𝒟_*it*_ = (*x*_*it*_, *n*_*it*_, *ϵ*_*it*_) represent the ALT allele counts, read depth, and sequencing error from the *t*^*th*^ time point. Thus, *x*_*i*1_, *x*_*i*2_, *x*_*i*3_ represent the allele counts from the first DNA sequence (2015), second DNA sequence (2022), and RNA sequence (2024), respectively. The likelihood of *θ*, the set of parameters comprising drift (*ψ, τ*) and mutation rate (*µ*), is expressed as ℒ (*θ*|𝒟_*i*_) ∝ ℙ(𝒟_*i*_|*θ*), where *ψ* = 1*/N*_*r*_ is the drift attributable to the transcriptional bottleneck and *τ* is the drift due to mtDNA turnover between 2015 and 2022. Dropping the heteroplasmy index *i* for clarity, we integrate ℙ(𝒟|*θ*) over the latent allele frequencies in the internal nodes of the tree:

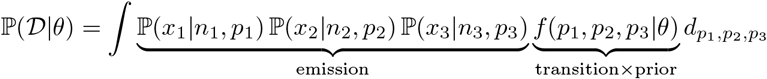

The observed allele counts are modeled as binomial samples from the true allele frequency at each time point accounting for the site-specific rate of sequencing error.

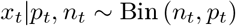

Because drift and mutation in different branches of the tree are independent, the joint distribution of allele frequencies, *f*(*p*_1_, *p*_2_, *p*_3_|*θ*) can be simplified into a product of conditionals:

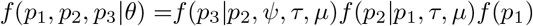

Note that the *p*_2_ → *p*_3_ transition is a function of both *ψ* (transcriptional bottleneck) and *τ* because the drift between 2022 and 2024 is modeled as 2*/*7 × *τ* (Fig. 2). The likelihood requires specifying *f*(*p*_1_), the distribution of allele frequency (DAF) representing uncertainty in the heteroplasmy fraction at the root of the tree. We model this using the posterior from a beta-binomial model with a flat prior such that *p*_1_|*x*_1_, *n*_1_ ∼ Beta(1 + *x*_1_, 1 + *n*_1_ − *x*_1_). This simplifies the likelihood to:

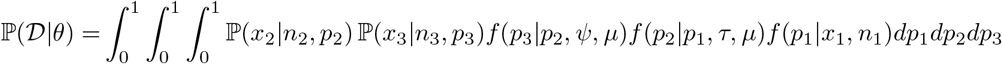

We model the evolution of heteroplasmies between internal nodes of the tree, i.e., the transition densities *f*(*p*_2_|*p*_1_, *τ, µ*) and *f*(*p*_3_|*p*_2_, *ψ, µ*) using an extension of the beta with spikes approximation of the Wright-Fisher model introduced in Tataru *et al*. (2015). Briefly, the DAF at node *t* follows a Wright-Fisher model where the mean and variance of *p*_*t*_ are functions of the allele frequency at the previous node (*p*_*t*−1_), mutation rate (*µ*), and the amount of drift (in scaled units of generations per effective population size *N*) accrued between the two nodes:

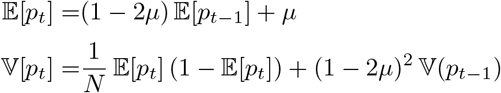

One can approximate the full DAF at *t* using the Beta distribution with shape parameters *α*_*t*_ and *β*_*t*_ which can be computed from the mean and variance in *p*_*t*_:

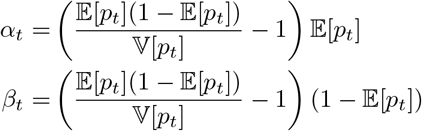

One limitation of the beta distribution is that it is defined for 0 *< p*_*t*_ *<* 1 and therefore, does not account for the possibility that alleles could be lost or fixed because of drift. The beta with spikes model accounts for this by adding point masses at *p*_*t*_ = 0 and *p*_*t*_ = 1:

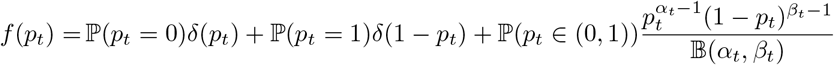

where 𝔹 and *δ* are the beta and Dirac delta functions, respectively, and ℙ(*p*_*t*_ ∈ (0, 1)) = 1 − ℙ(*p*_*t*_ = 0) − ℙ(*p*_*t*_ = 1). To fully specify *f*, we need the probabilities ℙ(*p*_*t*_ = 0) and ℙ(*p*_*t*_ = 1) of loss and fixation, respectively, and the shape parameters of the DAF conditional on the site being polymorphic. We can compute the conditional expectations and variance of *p*_*t*_|*p*_*t*_ ∈ (0, 1) using:

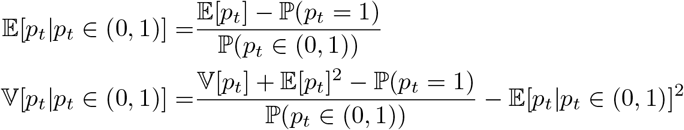

and then get the shape parameters *α*_*t*_ and *β*_*t*_ as shown above. To compute the loss and fixation probability in every generation, we assume that *µ* is small following Tataru *et al*., which results in the following approximations:

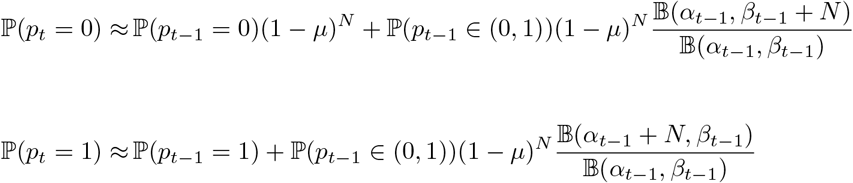

We compute the loss and fixation probabilities and the evolution of DAF recursively for each branch (of length *l*) by discretizing drift along the branch into generations of Δ = *l/N* where *N* = 1, 000 and computing *f*(*p*_*t*+Δ_|*p*_*t*_) as described above. The joint likelihood of observing all heteroplasmies is then computed assuming independence across sites:

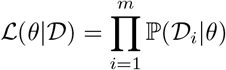

Our implementation differs from Tataru *et al*. in two ways. First, Tataru *et al*. use a single set of shape parameters to approximate the DAF across all sites at a given node, which can be imperfect when the number of sites is small, as in our case. To avoid this, we use a separate set of shape parameters for each site. Thus, the DAF represents the uncertainty in the heteroplasmy frequency at each site and we model the evolution of this uncertainty down the tree. Second, whereas Tataru *et al*. approximate the DAF on a discrete frequency grid, we represent it continuously using the beta-with-spikes distribution at each generation. Together, these extensions sacrifice runtime for higher resolution and accuracy. Our model accurately captures the behavior of heteroplasmy evolution under the Wright-Fisher model for a range of parameter values (Fig. S9).

We use a Bayesian approach to inference. Specifically, we estimate ℙ(*θ*|𝒟) ∝ ℒ (*θ*|𝒟) ℙ(*θ*) using differential evolution Markov Chain Monte Carlo (MCMC) implemented in the R package `BayesianTools` [57] using log-uniform priors on *ψ* ∈ [5 × 10^−4^, 0.5], *τ* ∈ [5 × 10^−4^, 0.5], and *µ* ∈ [10^−11^, 10^−8^]. We initialized ten chains with 50,000 iterations and a burn-in of 10,000. After visual inspection of the trace for convergence, we drew every 30th iteration to account for autocorrelation. We estimate the transcriptional bottleneck size — the *effective* number of copies of mtDNA that were transcribed — by assuming drift occurs in a single generation: 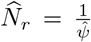. To validate our approach, we carried out simulations of heteroplasmy evolution under the Wright-Fisher model along the tree shown in Fig. 2A. To do this, we initialized 418 heteroplasmies at the root of the tree using allele frequencies from the real data, i.e., 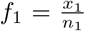. We used the Wright-Fisher model to simulate their evolution along the tree shown in Fig. 2A under a range of values for *τ* and *ψ* with an effective population size of 1,000 and mutation rate of 10^−7^.

### eQTL analysis

For *cis*-eQTL analysis, we merged the VCFs across all individuals using `bcftools merge` [58] with the -0 flag which replaces ./. genotypes with 0/0. For *trans-*eQTL analysis, we used the genotypes at 1.06M SNPs from the HapMap3 SNP list. We carried out *cis-*eQTL analysis of mtDNA gene expression in R using a linear model where we included sex, 5 gPCs, batch, and 100 ePCs as covariates. To correct for multiple testing, we first computed the effective number of genes (*m*_*eff*_) through eigendecomposition of the mtDNA gene expression correlation matrix. Specifically, *m*_*eff*_ is defined as the minimum number of PCs required to explain >99% of the expression variance (≈ 10). This resulted in a study-wide p-value threshold of 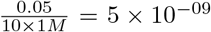. *Trans*-eQTL analysis was done using `TensorQTL` [59]. Fine-mapping was done with `SuSiE` [22] using LD matrices generated from genotypes residualized on the first five gPCs.

To understand how eQTL discovery relates to the selective constraint on a gene and its expression level, we downloaded nuclear constraint scores from gnomAD (v4.1, 807,162 individuals, released on 04/19/2024) [29] and mtDNA constraint scores from Lake *et al*. [28]. We defined constraint as the ratio between the observed and expected number of missense mutations such that a lower score implies more constraint. We chose missense mutations as opposed to loss-of-function (LOF) mutations to keep constraint comparable between the nuclear and mitochondrial genomes, for which the LOF constraint was not reported [28]. To account for statistical noise, we chose the upper bound of the 90% confidence interval of the observed vs expected ratio. We categorized the 10% of genes with the lowest score as ‘highly constrained’ and the remaining 90% as ‘less constrained’. We modeled the effect of gene expression and constraint on eQTL discovery in MAGE with a quasipoisson generalized linear model implemented with the `glm()` function in R with number of fine-mapped credible sets for each gene as response and the mean TMM and constraint category as predictors.

## Data and Code Availability

The MAGE data is publicly available from Sequence Read Archive under accession PRJNA851328. The 1000 Genomes Project low-coverage and high-coverage sequencing is publicly available at https://www.internationalgenome.org/data-portal/data-collection/30x-grch38. Expression count matrices and covariates from GTEx v8 are available at https://gtexportal.org/home/downloads/adult-gtex/bulk_tissue_expression. GTEx Protected Data (e.g. whole-genome alignments and genotypes) are available under dbGaP Study Accession phs000424.v10.p2 and were analyzed under project number 34331. Full summary statistics for all association analyses and *trans*-eQTL mapping are available on Zenodo (https://doi.org/10.5281/zenodo.20350769 [60]). All code is freely available at the following GitHub repository https://github.com/zaidilab/mtDNA_GE.

## Acknowledgements

Thanks to members of the Zaidi Lab, Frank Albert, members of the Albert Lab, and Kateryna Makova for helpful discussions and feedback. Thanks to the Biogen Administrative Center and Minnesota Supercomputing Institute at the University of Minnesota for their support. This research was funded by NIH/NIGMS awards R00GM137076 and R35GM159945 to AAZ and NIH award T32GM140936 to STB. The content is solely the responsibility of the authors and does not necessarily represent the official views of the National Institutes of Health.

## Author Contributions

Conceptualization: AAZ; Data curation: DT, RM, PR, SR, BL, SR, AAZ; Formal analysis: PR, SR, BL, STB, SR, AAZ; Funding acquisition: AAZ; Methodology: AAZ, Supervision: SR, AAZ; Visualization: PR, SR, BL, SR, AAZ; Writing – original draft: PR, BL, AAZ; Writing – review and editing: STB, RM, SR, AAZ.

## Supplemental Information

Document S1. Figures S1-S13

Data S1. mtDNA copy number estimates from high- and low-coverage WGS data for MAGE individuals. Data S2. Variant allele counts and sequencing depth for 418 heteroplasmies detected with VAF > 0.01 and depth > 1000× in the low-coverage WGS data of the MAGE individuals. Data S3. Full *cis*-eQTL result of mtDNA gene expression in MAGE. Data S4. Fine-mapping results of *cis*-eQTL analysis of mtDNA gene expression in MAGE. Data S5. Fine-mapping results of *trans*-eQTL analysis of mtDNA gene expression in MAGE.

## Supplementary Material

**Figure S1:**
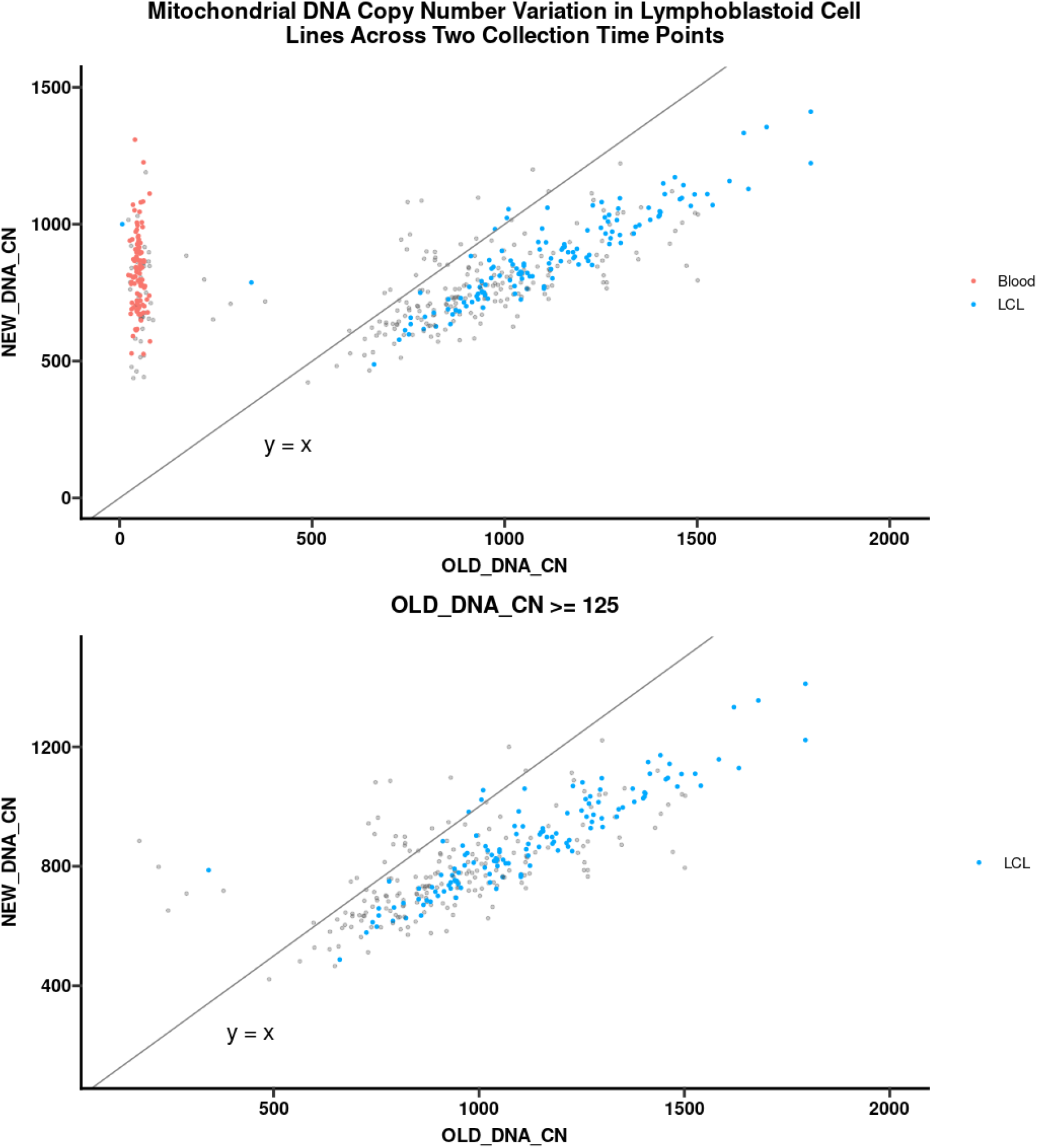
Comparison between mtDNA copy number estimated from low-coverage [16] (x-axis) and high-coverage [15] (y-axis) sequencing of the 1000 Genomes Project. Each point is an individual and the colors represent the tissue from which the sample was drawn for low-coverage sequencing. Grey indicates that tissue information was not available. The high-coverage sequencing was carried out from lymphoblastoid cell lines (LCLs). The *y* = *x* line is also shown. The correlation is shown for (A) all samples or (B) only samples from the low-coverage effort predicted to be sequenced from LCLs.

**Figure S2:**
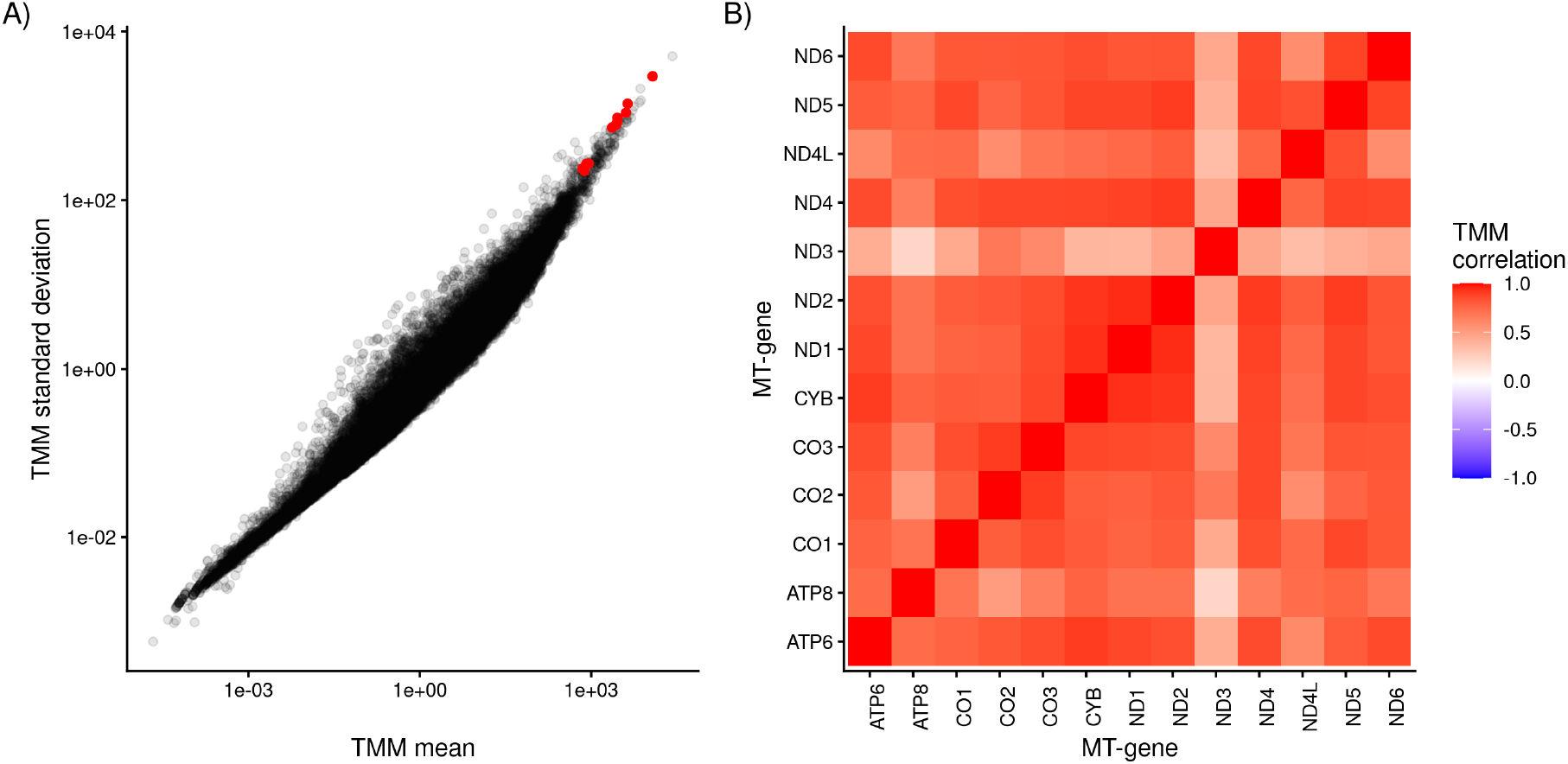
(A) Mean (x-axis) and standard deviation (y-axis) of protein-coding genes in MAGE. MtDNA-encoded genes are shown in red. (B) Correlation in gene expression between protein-coding genes encoded by the mtDNA.

**Figure S3:**
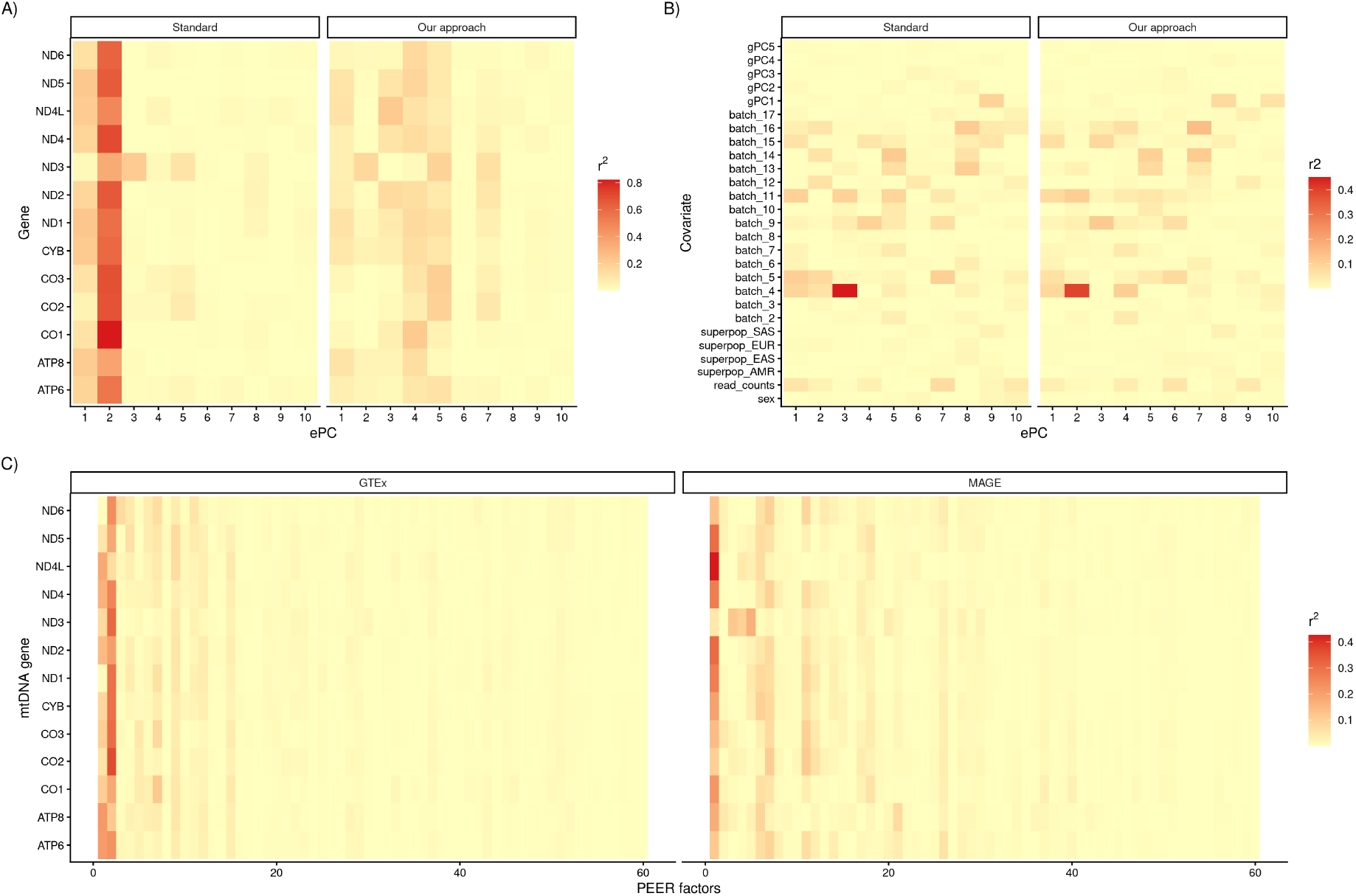
Heatmap showing the coefficient of determination (*r*^2^) (A) between ePCs and mtDNA-encoded gene expression, (B) between ePCs and commonly used covariates, and (C) between PEER factors and mtDNA gene expression. In (A) and (B), the facets represent the set of genes used to normalize gene expression and compute ePC. “Standard” represents all protein coding genes and “Our approach” represents protein coding genes excluding mtDNA-encoded and nuclear-encoded mitochondrial genes. Top ePCs computed with our approach are not as strongly correlated with expression of mtDNA genes as the standard approach. Top ePCs computed with our approach still retain information about important biological and technical covariates. (C) shows the *r*^2^ between PEER factors computed by GTEx [18] (Left) and MAGE [12] (Right) and mtDNA-encoded gene expression.

**Figure S4:**
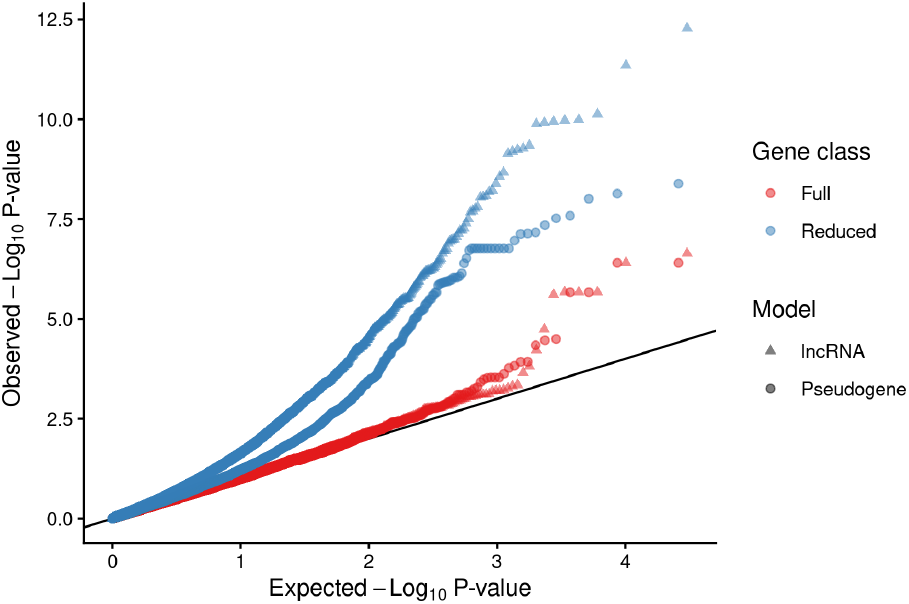
QQplot of association between mtCN and expression of lncRNA (triangles) and pseudogenes (points) in MAGE. Results are shown for two models, one with sex, age, age^2^, batch, and gPCs (reduced), and second model (full), which also includes ePCs. Pseudogenes and lncRNAs were both excluded from the normalization and ePC calculation.

**Figure S5:**
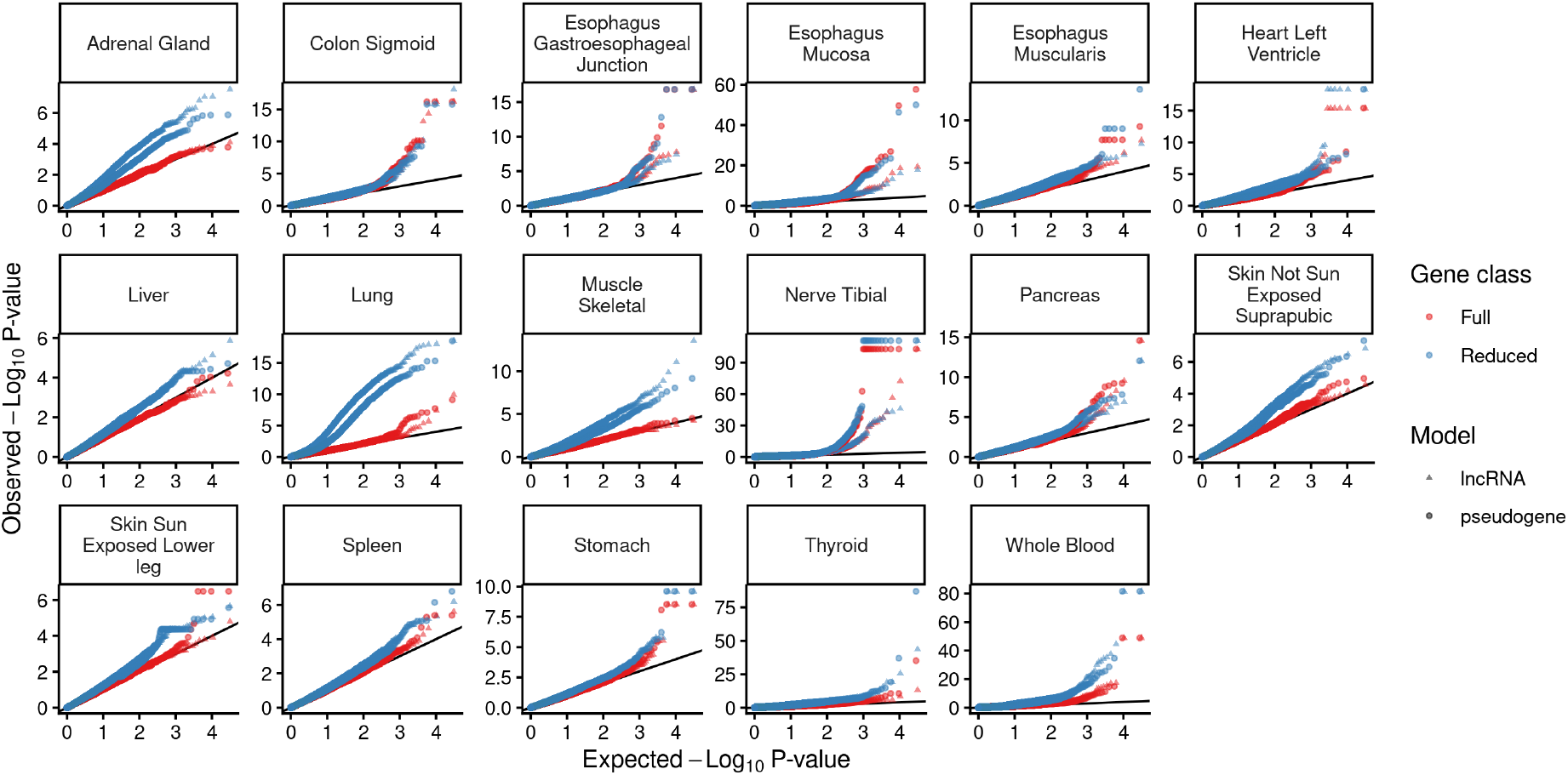
QQplot of association between mtCN and expression of lncRNA (triangles) and pseudogenes (points) in GTEx. Only tissues where the number of samples were greater than the number of model covariates are shown. Results are shown for two models, one with sex, age, age^2^, batch, and gPCs (reduced), and second model (full), which also includes ePCs.

**Figure S6:**
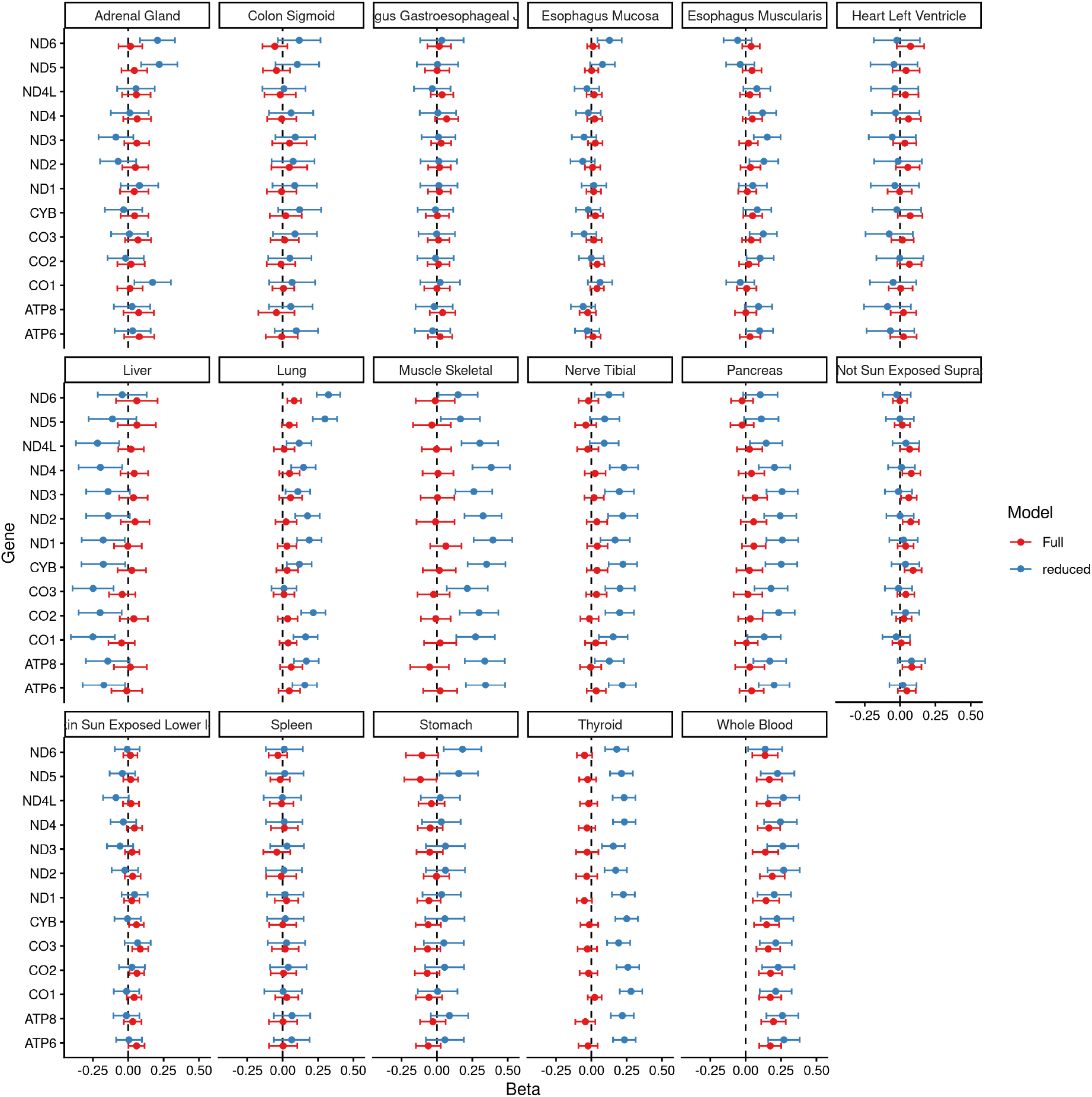
Association between mtCN and expression of mtDNA-encoded protein-coding genes (y-axis) in GTEx. The effect sizes (x-axis) are standardized. Results are shown for two models, one with sex, age, age^2^, batch, and gPCs (reduced), and second model (full), which also includes ePCs.

**Figure S7:**
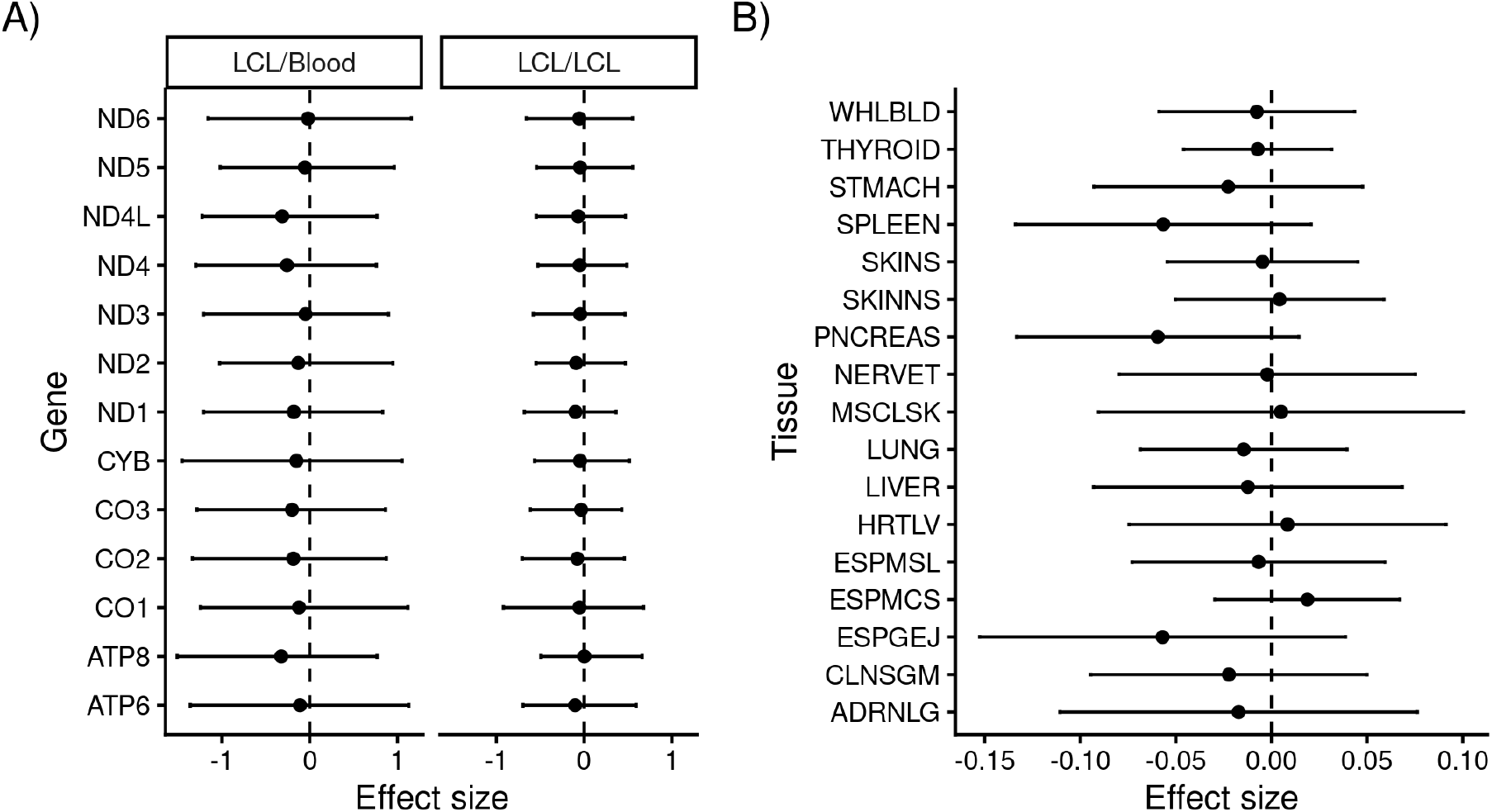
Association between predicted mtCN (pmtCN) and mtDNA gene expression in (A) MAGE and (B) GTEx. (A) The x-axis shows the estimated effect of pmtCN on expression of 13 protein-coding mtDNA genes (y-axis) in MAGE. The pmtCN was either constructed from effects estimated in the original GWAS, which was conducted in whole blood (left) or effects re-estimated in LCLs (right) (Methods). (B) Shows the same effect only for the COX1 mtDNA gene only in GTEx tissues (y-axis).

**Figure S8:**
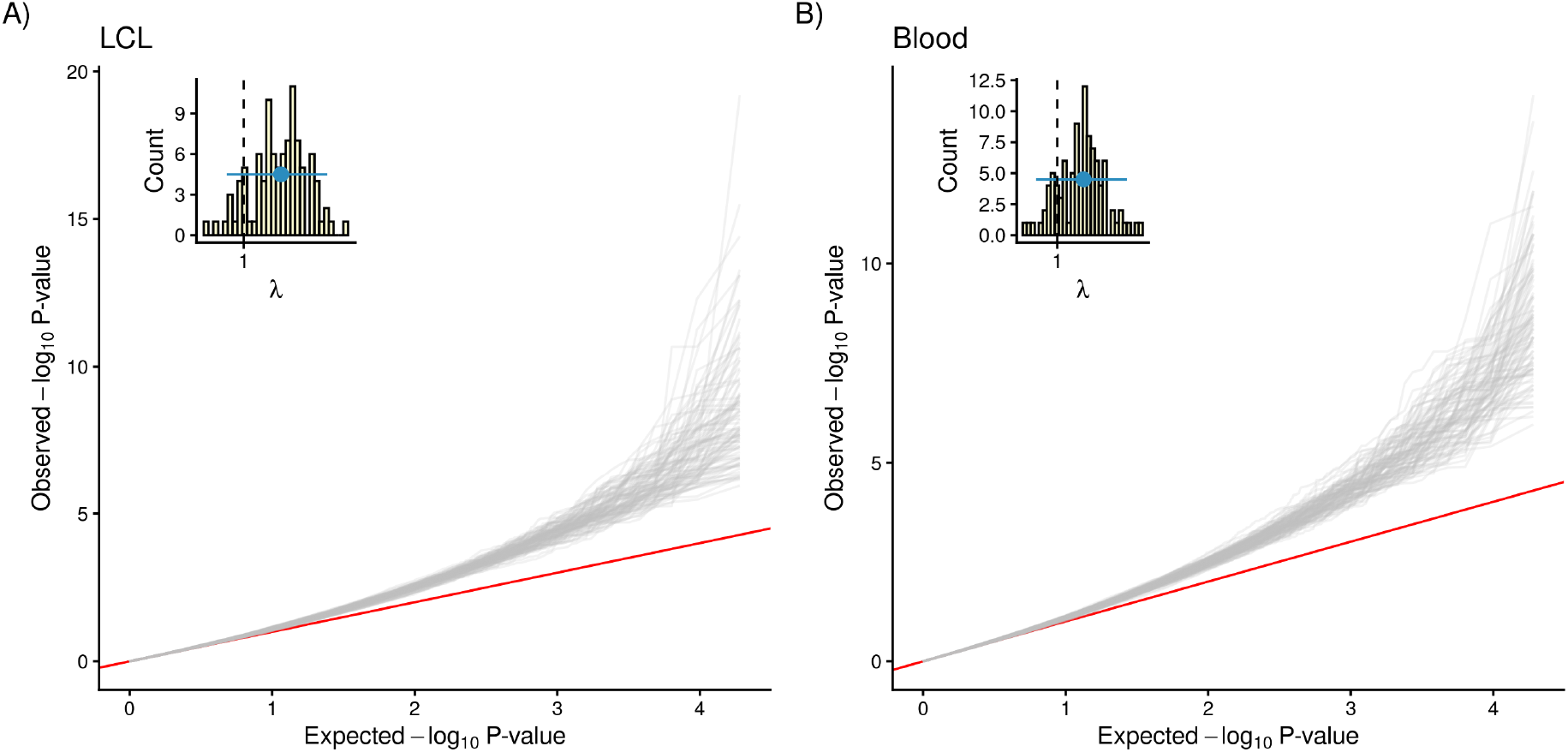
QQplots of the association between predicted mtCN (pmtCN) and pseudogene expression in MAGE. The loci used to compute pmtCN were selected from a previous GWAS[47]. The effect sizes used to compute pmtCN were either (B) taken directly from the previous GWAS [47], which was done in whole blood or (A) re-estimated in an independent subset of lymphoblastoid cell lines (LCLs) in MAGE (Methods). Each grey line represents a bootstrapped sample and the distribution of corresponding inflation factors (*λ*) are shown in the inset.

**Figure S9:**
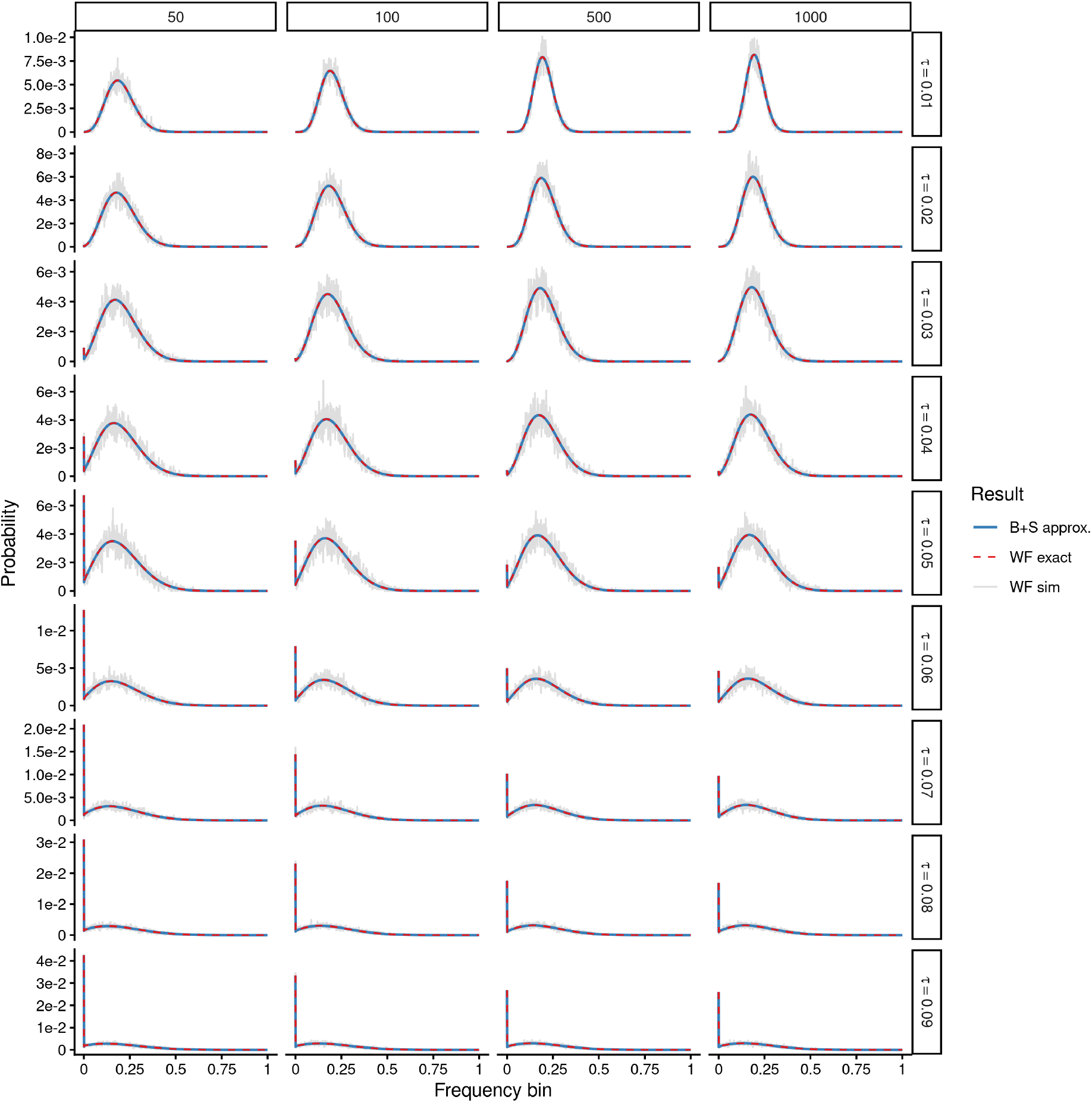
Our model accurately captures heteroplasmy evolution under a range of *τ* (rows) and *N*_*r*_ (columns) parameters. Each panel shows the distribution of allele frequency (DAF) of a heteroplasmy initialized at a frequency of 0.2. Grey lines represent simulated DAF (10,000 replicates) and the red dashed and solid blue lines represents the theoretical expectation under the exact Wright-Fisher model and beta with spikes approximation.

**Figure S10:**
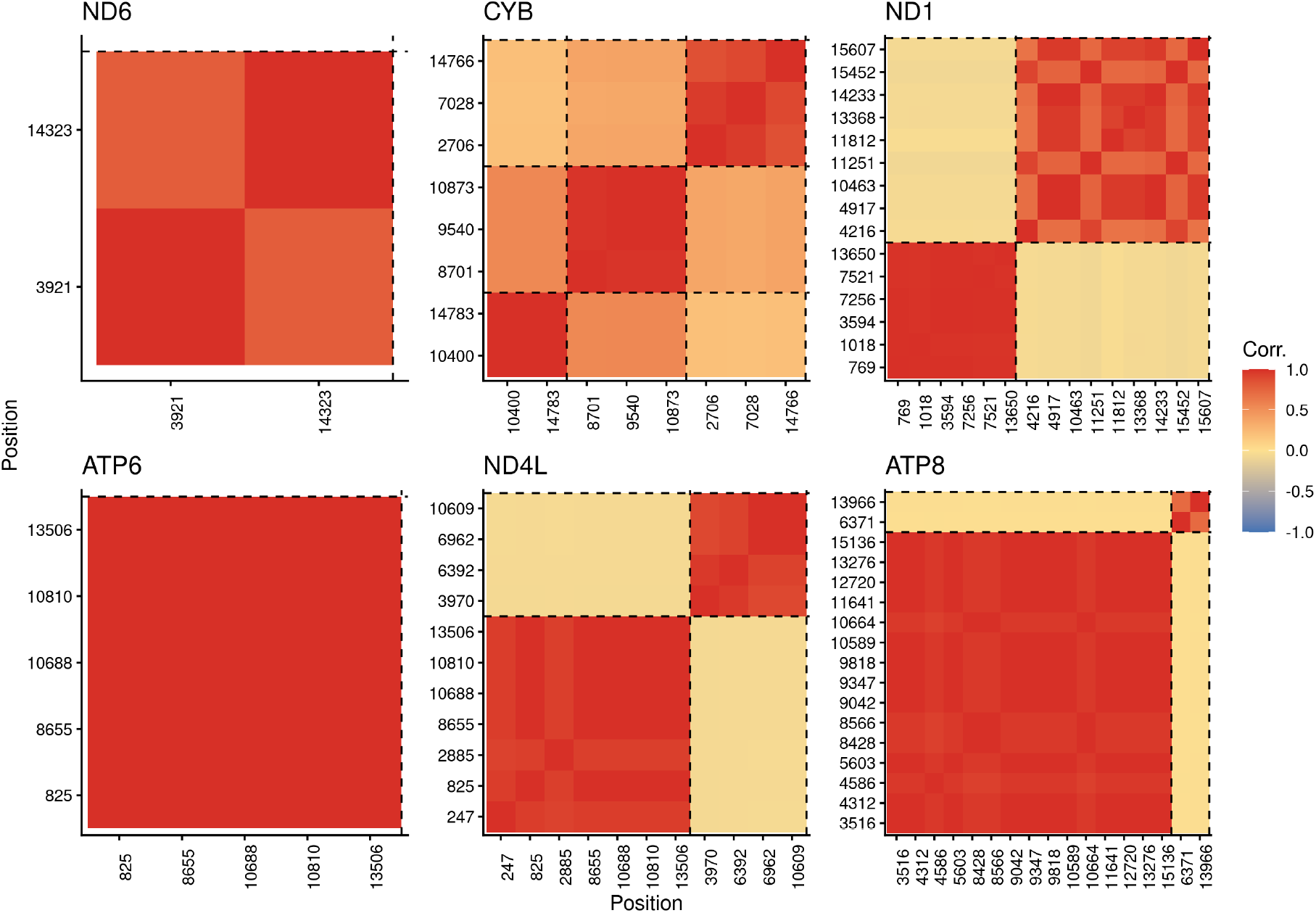
LD across SNPs in fine-mapped credible sets. For each mtDNA gene, SNPs are grouped first by credible set, which are separated by dashed lines, and sorted by position.

**Figure S11:**
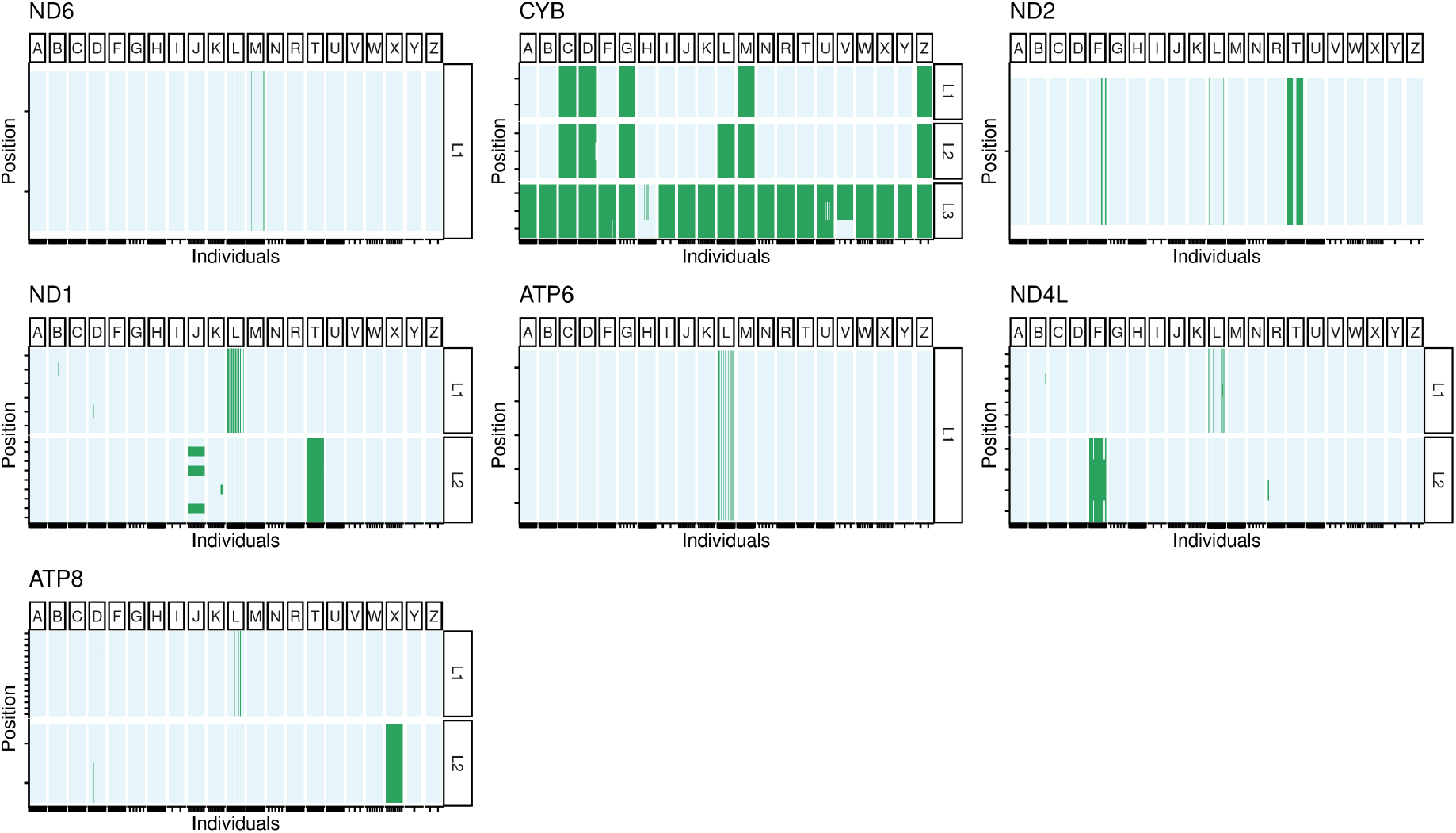
Haplotype structure of SNPs in fine-mapped credible sets. Reference alleles are shown in blue and alternate alleles in green. Individuals (x-axis) are split by major haplogroup and SNPs (y-axis) by credible set. Ticks on either axes represent the density of data points in each facet.

**Figure S12:**
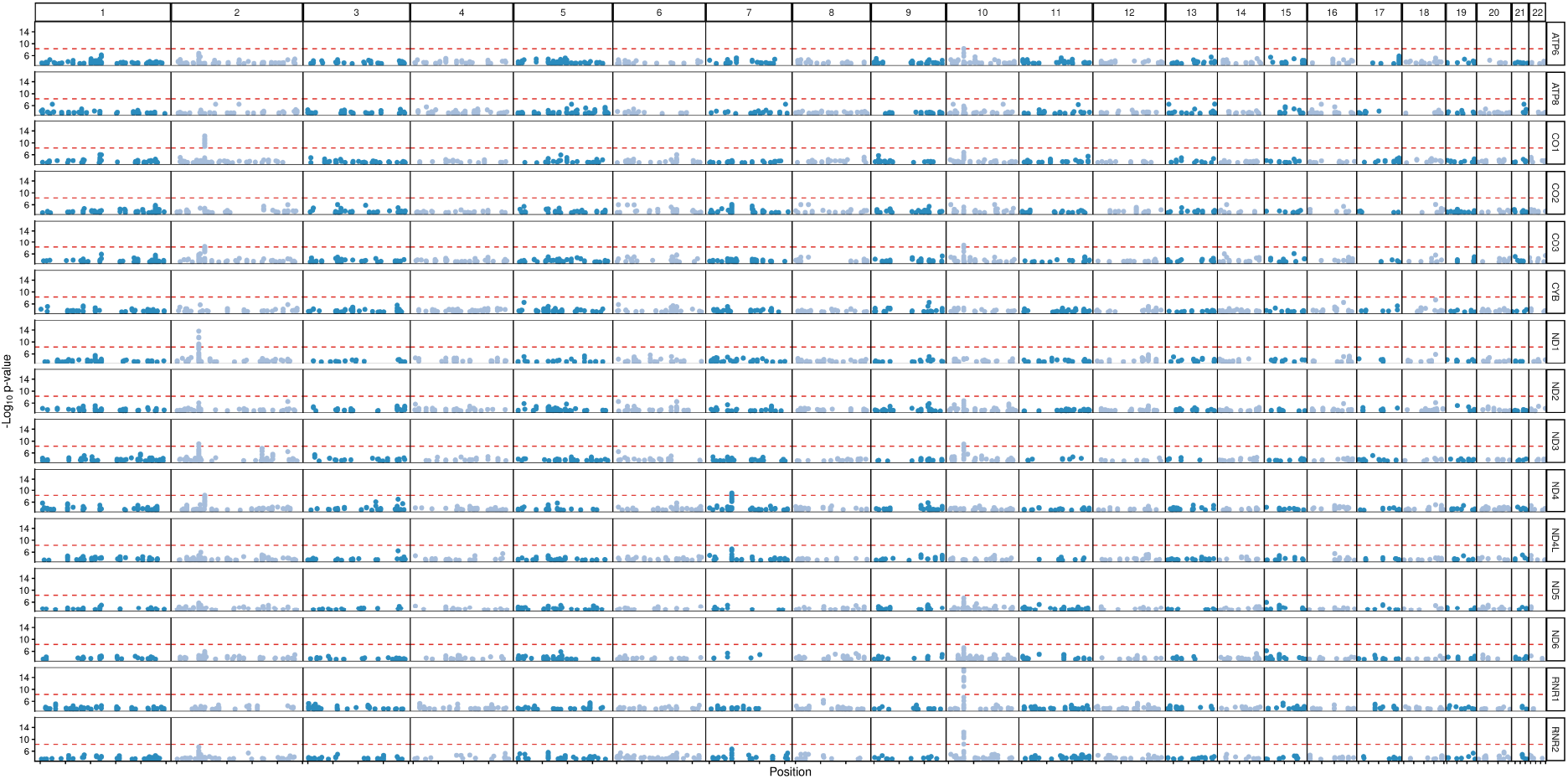
Manhattan plot of *trans*-eQTL analysis of expression of mtDNA genes (rows) in MAGE. Only SNPs with a p-value < 5 × 10^−04^ are shown for simplicity. The red horizontal dashed line shows the study-wide threshold of 5 × 10^−09^.

**Figure S13:**
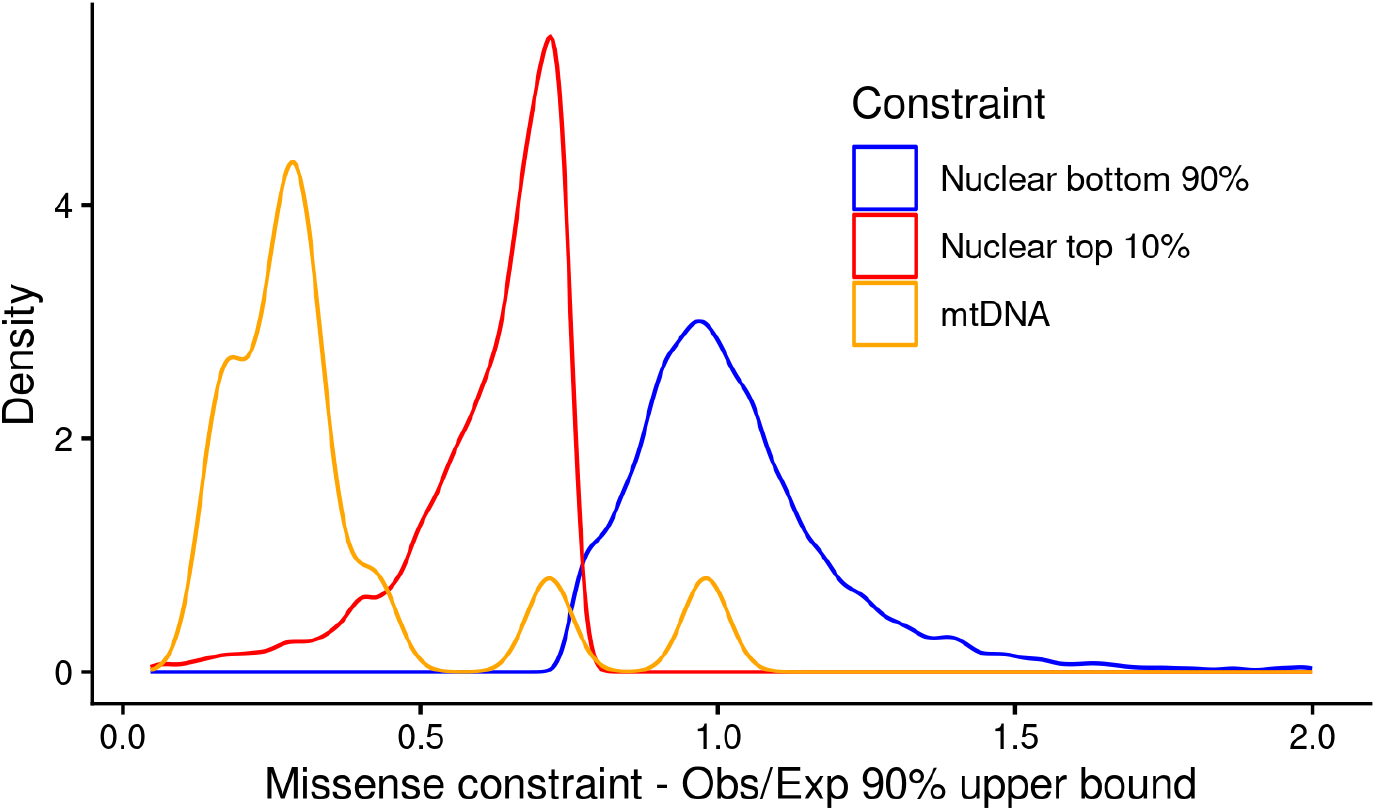
Distribution of selective constraint (upper bound of the 90% CI of observed/expected missense variants) for nuclear genes in the top 10% (red) and bottom 90% (blue) in terms of constraint. The missense constraint scores for mtDNA protein-coding genes are shown in orange.

